# Functional and comparative genomics reveals conserved noncoding sequences in the nitrogen-fixing clade

**DOI:** 10.1101/2021.07.27.453985

**Authors:** Wendell J. Pereira, Sara Knaack, Daniel Conde, Sanhita Chakraborty, Ryan A. Folk, Paolo M. Triozzi, Kelly M. Balmant, Christopher Dervinis, Henry W. Schmidt, Jean-Michel Ané, Sushmita Roy, Matias Kirst

## Abstract

Nitrogen is one of the most inaccessible plant nutrients, but certain species have overcome this limitation by establishing symbiotic interactions with nitrogen-fixing bacteria in the root nodule. This root nodule symbiosis (RNS) is restricted to species within a single clade of angiosperms, suggesting a critical evolutionary event at the base of this clade, which has not yet been determined. While genes implicated in the RNS are present in most plant species (nodulating or not), gene sequence conservation alone does not imply functional conservation – developmental or phenotypic differences can arise from variation in the regulation of transcription. To identify putative regulatory sequences implicated in the evolution of RNS, we aligned the genomes of 25 species capable of nodulation. We detected 3,091 conserved noncoding sequences (CNS) in the nitrogen-fixing clade that are absent from outgroup species. Functional analysis revealed that chromatin accessibility of 452 CNS significantly correlates with the differential regulation of genes responding to lipo-chitooligosaccharides in *Medicago truncatula*. These included 38 CNS in proximity to 19 known genes involved in RNS. Five such regions are upstream of *MtCRE1*, *Cytokinin Response Element 1,* required to activate a suite of downstream transcription factors necessary for nodulation in *M. truncatula*. Genetic complementation of a *Mtcre1* mutant showed a significant association between nodulation and the presence of these CNS, when they are driving the expression of a functional copy of *MtCRE1*. Conserved noncoding sequences, therefore, may be required for the regulation of genes controlling the root nodule symbiosis in *M. truncatula*.

## INTRODUCTION

Nitrogen is one of the most limiting plant nutrients, despite making up 78% of the Earth’s atmosphere. Plants cannot access nitrogen directly and must obtain it from the soil. Some species have evolved to overcome this limitation by establishing symbiotic associations with nitrogen-fixing bacteria. In some of these mutually beneficial relationships, the host plant develops root nodules, new root organs that provide an environment for the bacteria to fix nitrogen. Root-nodule symbioses (RNS) evolved from the recruitment of molecular and cellular mechanisms from the arbuscular mycorrhizal symbiosis and the lateral root developmental program (Gualtieri and Bisseling 2000; Streng et al. 2011; Schiessl et al. 2019). Still, the specific molecular origins of the trait remain unknown.

Plants able to develop RNS are restricted to species in a single monophyletic clade of angiosperms, consisting of the orders Rosales, Fagales, Cucurbitales, and Fabales (Soltis et al. 1995). While this nitrogen-fixing clade (NFC) comprises 28 families, only ten include species capable of developing nodules. Within the majority of these families, most of the member species lack this capability. The scattered distribution of the RNS across the NFC suggests it either evolved independently multiple times (convergent evolution) or was lost after one or more gain events. Recent work identified essential genes specific to the RNS such as, *NODULE INCEPTION* (*NIN*), *RHIZOBIUM-DIRECTED POLAR GROWTH* (*RPG*) and *NOD FACTOR PERCEPTION* (*NFP*), and the loss of either is associated with the absence of the trait in the NFC (Griesmann et al. 2018; Velzen et al. 2018). While the loss of these genes may partially explain the evolution of symbiosis in this clade, their presence in species outside of the NFC indicates that other genetic factors are likely to underlie the trait origin. Moreover, the observation that nodulating species are confined to a specific clade provides evidence for a single origin or evolutionary event behind the genetic underpinning nodulation at the NFC base (Soltis et al. 1995; Werner et al. 2014). The nature of this event has not been determined. However, its discovery is a critical step towards understanding the evolutionary emergence of nodulation and its engineering into crops, a long-term goal for sustainable agriculture.

While genes implicated in nodulation are present in most species (nodulating or not) within the NFC and even outside the clade, gene sequence conservation alone does not imply maintenance of similar function. Closely related species can exhibit extensive developmental or phenotypic divergence, despite having high gene content similarity (Kirst et al. 2003). Such differences are often due to variation in the transcription regulation (Pollard et al. 2006; Prabhakar et al. 2006). Gene expression regulation is in part determined by the interaction of transcription activators and repressors with specific regulatory sequences. Epigenetic factors, such as the accessibility of the genomic regions containing these regulatory sequences, further modulate their contribution to gene expression.

Species with the ability to establish symbiosis with nitrogen-fixing bacteria are expected to share conserved sequences due to negative (purifying) selection, including conserved noncoding sequences (CNS). These CNS often carry functionally critical phylogenetic footprints and harbor transcription factor binding motifs (TFBM) or other *cis*-acting binding sites (Freeling and Subramaniam 2009). They can be required to maintain the conformational structure of mRNAs, and, when located in introns, they may be necessary for correct gene splicing. Conserved noncoding regions have been extensively documented in plants (Haudry et al. 2013; Burgess and Freeling 2014; de Velde et al. 2014; Liang et al. 2018) and shown to perform essential roles in regulatory networks and plant development. In maize, more than half of putative *cis*-regulatory sequences overlaps with CNS. Similarly, TFBMs, eQTLs and open chromatin regions are enriched in these noncoding sequences (Song et al. 2021). In Arabidopsis, CNS overlaps with a large fraction of open chromatin sites and functional TFBMs (de Velde et al. 2014). Therefore, regulatory sequences critical to RNS may be identified by searching for CNS in the genomes of nodulating species in the NFC. Furthermore, chromatin accessibility of these regions may also be associated with the developmental process of nodulation, contributing to the regulation of gene expression.

Orthologous CNS that are more similar to each other than expected from neutral evolution, and accessible during nodulation, may have critical functional roles in this process. Moreover, the identification of CNS among nodulating species in the NFC but diverged in species outside of the clade could point to regulatory sequences that underlie the origin of the nodulation trait. Here we pursued the identification of putative regulatory sequences associated with the origin of the RNS based on the hypothesis that a genetic event (predisposition or gain) occurred at the NFC base. We identified several CNS potentially involved in the regulation of genes that are required for nitrogen-fixation. Furthermore, we demonstrate that the deletion of such regions upstream of *MtCRE1* significantly reduces the number of nodules produced by *M. truncatula* when symbiosis is established with *Sinorhizobium meliloti*. Such regulatory sequences may have contributed to the differential regulation of genes critical for nodule development and, consequently, be important for engineering nodulation in crops outside of the NFC.

## RESULTS

### An atlas of conserved non-coding sequences in the nitrogen-fixing clade

#### Selection of genomes of species in the NFC and whole-genome alignments

We identified genome sequences from 84 species in the NFC orders Fabales, Fagales, Cucurbitales, and Rosales in NCBI. After removing those species unable to associate with nitrogen-fixing bacteria, a total of 33 genomes remained. Assessment of assembly contiguity and gene completeness resulted in the exclusion of eight additional genome sequences. Thus, the genomes of 25 species capable of RNS, including the reference *M. truncatula*, were used to identify CNS (Figure 1; Supplemental File S1). Moreover, we selected a group of nine species belonging to orders Brassicales, Linales, Malpighiales (three species), Malvales, Myrtales, Sapindales, and Vitales (all outside the NFC; Figure 1) to be used as an outgroup (see Methods).

**Figure 1.**
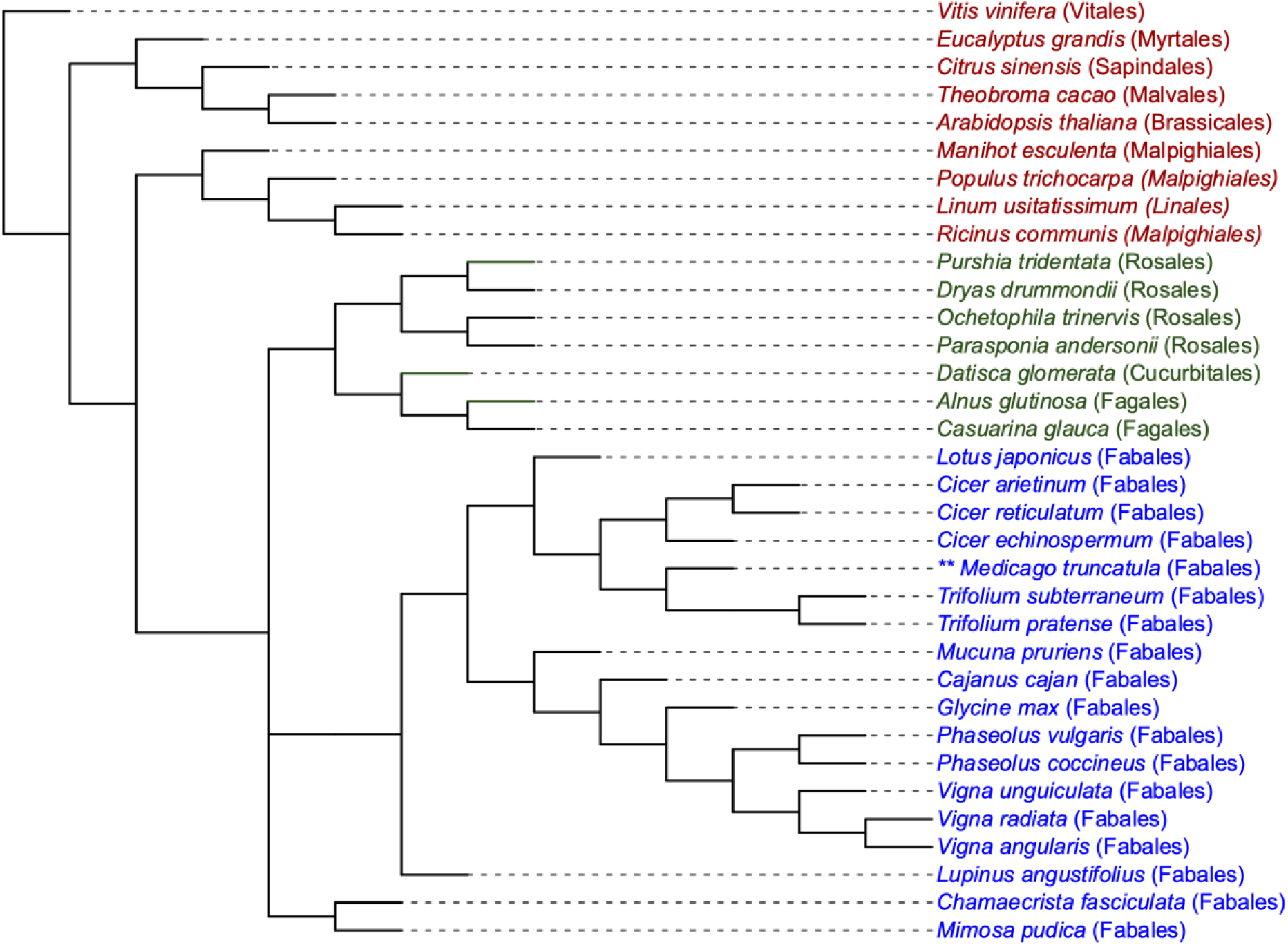
Phylogenetic tree used to guide the creation of the multiple genome alignment by ROAST. Presented are 25 species included in the evolutionary analysis, belonging to the NFC clade (capable of nitrogen fixation) orders Fabales (blue), Fagales, Cucurbitales and Rosales (green). Additionally, nine outgroup species (red) were included. Asterisks indicate the species used as reference to produce the whole-genome alignments.

The pairwise whole-genome alignment of each species to *M. truncatula* covered on average 16.2% (s.d. = 0.06) of nucleotides of the reference genome and 15.0% (s.d. = 0.06) of nucleotides in the genomes of the remaining species in the NFC. In the pairwise whole-genome alignments of the outgroup’s species, 8.6% (s.d. = 0.01) of nucleotides in the *M. truncatula* and 10.7% (s.d. = 0.05) of the nucleotides of the other species were covered, on average (Supplemental File S1). As expected, a higher degree of alignment coverage was detected for the species in the NFC clade, which are more phylogenetically related to *M. truncatula*. Moreover, a large share of the regions covered by the alignment of the NFC group corresponds to coding sequences in *M. truncatula*. On average, 40.9% (s.d. = 11.6) of the nucleotides of alignments between the reference and target species were located within these regions. Considering the coding sequences in the *M. truncatula* genome, an average of 56.4% (s.d. = 0.08) of the nucleotides were covered in the alignment of the NFC group, and 45% (s.d. = 0.03) in the outgroup (Supplemental File S1). For each of these two groups, multiple whole-genome alignments were generated using a phylogenetic guided approach and used to search for conserved regions.

#### Identification of conserved sequences

We used PhastCons to estimate the conservation score across the genome of species in the NFC. PhastCons identifies conserved sequences in multiple genome alignments while considering the phylogenetic relationship and nucleotide substitution in each site under a neutral evolutionary model. PhastCons detected 114,173 conserved regions (mean of length = 96.63 bp, s.d. = 107.7) from the alignments of 25 NFC species, with a conservation score higher than 0.5 and a size of five or more base pairs (Table 1). The conserved regions represented 2.5% of the *M. truncatula* genome. Next, we also applied PhastCons to identify conserved regions among species in the outgroup and detected 97,302 such regions (mean of length = 104.63 bp, s.d. = 104.41).

**Table 1.** Descriptive statistics of the length of all conserved regions identified by PhastCons and of the set of regions considered as true CNS.

| **Group** | **N° of**<br>**regions** | **Min**<br>**(bp)** | **Max**<br>**(bp)** | **Mean**<br>**(bp)** | **Median**<br>**(bp)** | **S.D.**<br>**(mean)** | **C.I. (95%)** |
| --- | --- | --- | --- | --- | --- | --- | --- |
| **Conserved regions**<br>**(PhastCons)** | 114,173 | 5 | 3,111 | 96.63 | 70 | 107.7 | 93.01 – 94.26 |
| **CNS** | 5,199 | 5 | 431 | 43.04 | 27 | 43.57 | 41.86 – 44.22 |

The conserved regions identified by PhastCons were mainly located within genic regions or their vicinity. Among the 114,173 conserved regions, 107,444 were within (overlap of one or more nucleotides) a coding sequence. For the remaining conserved regions, 1,526 were located downstream (0-2kb), 485 long-range (distal) downstream (2-10kb), 2,924 upstream (0-2kb), and 537 long-range (distal) upstream (2-10kb) of a gene. Moreover, 1,093 of these regions were within introns and 164 in intergenic regions, further than 10kb of the transcription start or transcription stop site of any gene.

#### Selection of CNS within the NFC and their genomic context

To focus on the conserved noncoding sequences, we excluded from the further analysis the majority (>94%) of the 114,173 regions detected by PhastCons, which overlapped with coding sequences. After removing the conserved regions within coding sequences, 6,729 (5.9%) remained. The CNS located outside of coding sequences were further investigated to determine if they represent noncoding sequences such as noncoding RNAs, transposable elements (TEs), or coding genes not described in the *M. truncatula* genome annotation. The abundance of these noncoding regions in plant genomes may result in the detection of high PhastCons conservation scores, while being unrelated to the evolutionary importance of RNS.

Among the 6,729 conserved regions located outside coding sequences, 515 and 778 were excluded from further analysis due to overlap with noncoding RNAs and TEs, respectively. Moreover, 225 conserved regions had significant similarity (blastx, e-value < 0.01) to sequences in the non-redundant protein database and were removed. These regions may be contained within pseudogenes or coding sequences missing from the current genome annotation of *M. truncatula*. Twelve regions were removed because they were from mitochondrial or chloroplast genomes. After filtering for the criteria described above, 5,199 conserved regions were considered *bona fide* CNS. These represent 4.6% of all conserved regions identified by PhastCons. The length of these CNS was significantly smaller than the observed for the complete set of conserved regions identified by PhastCons (Table 1; Supplemental Fig S1). This observation possibly reflects shorter regulatory motifs detected in noncoding sequences compared to coding sequences.

The majority of the 5,199 CNS (2,441 or 46.95%) were detected upstream of the translation start site (TSS) of the closest gene (0-2 kb), based on the *M. truncatula* reference genome. Among the remaining CNS, 1,245 (23.95%) were downstream of the translation end site (0-2 kb) and 871 (16.75%) in introns. An additional 294 (5.65%) CNS were located between two and ten kilobases upstream the transcription start site and were classified as distal upstream. Also, 275 CNS were located between two and ten kilobases downstream the transcription stop site and were classified as distal downstream. All other CNS (73) were found in intergenic regions (Supplemental Fig S2).

### Orthologous CNS in *M. truncatula* and *G. max*

For a CNS to be evolutionarily and biologically relevant across taxa, it is expected to occur in a similar genomic context (for instance, within the promoter region of orthologous genes). Thus, we next examined if the CNS identified in the *M. truncatula* genome were in orthologous regions of the soybean genome, *Glycine max*; v.2.1 (Schmutz et al. 2010). *Glycine max* was selected for this analysis because gene orthology relative to *M. truncatula* is well-established and described in the PLAZA database (Van Bel et al. 2018). From the 5,199 CNS, it was possible to recover the coordinates of 5,165 in the *G. max* genome. The distances of these CNS relative to the closest gene were calculated and classified as described above for *M. truncatula*. Most of the CNS were classified similarly in both genomes (4,098 CNS; 78.82%), especially in the upstream, downstream, and intronic categories (Figure 2, Supplemental File 2).

**Figure 2.**
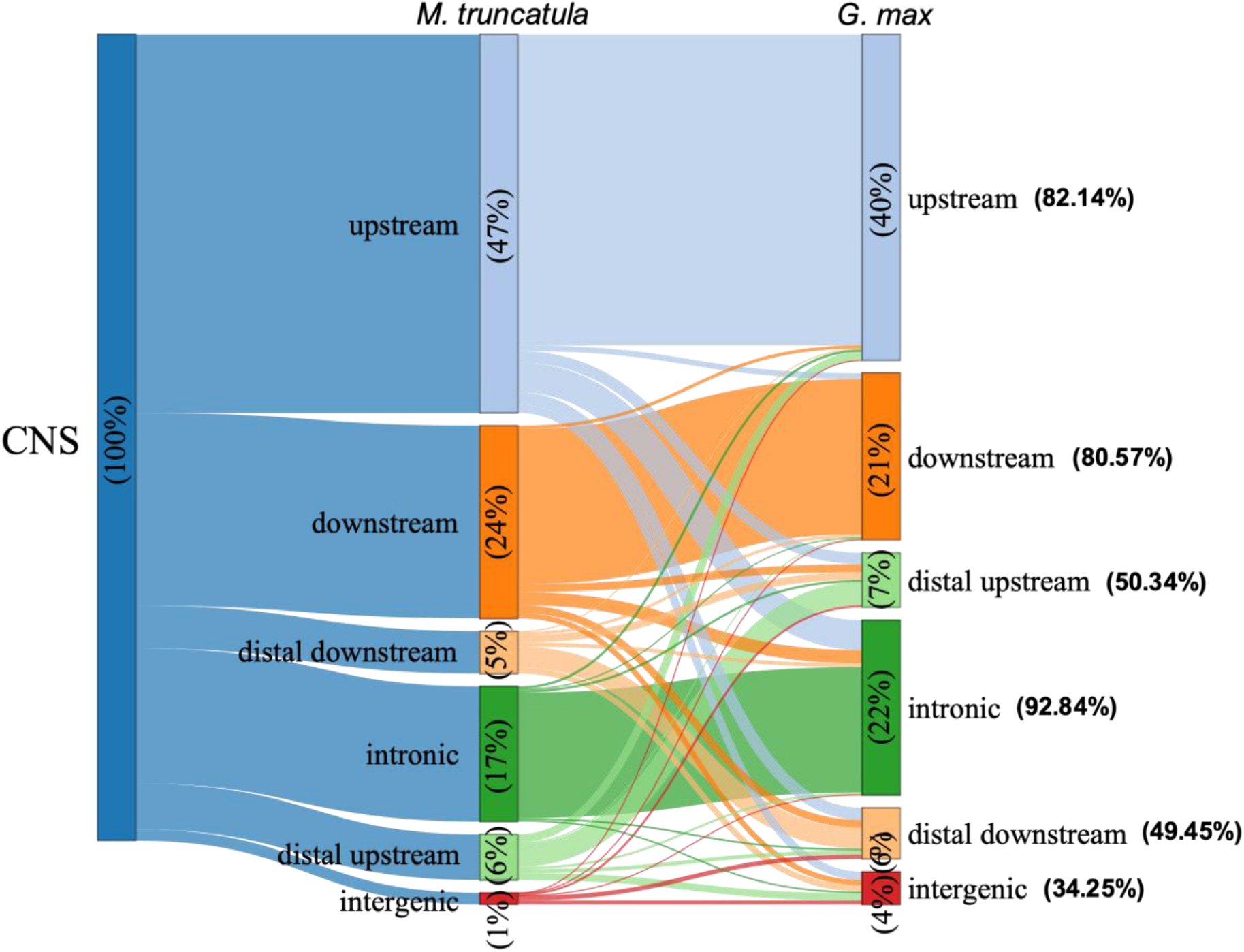
Classification of CNS according to their distance to the closest genes in the genomes of *M. truncatula* and *G. max*. The percentages within the bars represents the distribution of the CNS in each species among the six categories. On the right, the percentage of CNS that have the same classification in *G. max* as in *M. truncatula* is shown.

The most prominent disagreement was observed for the CNS classified as intergenic, where only 34.25% of those in this category in *M. truncatula* were in the same category in *G. max*. Distal upstream and distal downstream also presented a significant disagreement in their classification. Approximately 50% of the CNS were classified equally in both genomes for these two categories (Figure 2). This lack of agreement is expected since conserved noncoding sequences are, in general, closely located to their target genes, and CNS found further than 2 kb from genes may not be functionally relevant. Nonetheless, long-range *cis*-regulatory elements are also known in plants and the distal CNS represent a potential approach to their detection.

Next, we assessed if the closest gene to each CNS classified in the same category (except for intergenic CNS) in both *M. truncatula* and *G. max* genomes correspond to orthologous genes in these species. For several of the CNS it was not possible to determine if the closest genes in both species were orthologous. This happened because there was no correspondence between *M. truncatula* v4 and v5 gene IDs, or between the Entrez ID and the *G. max* IDs used by PLAZA, or yet no orthology mapping established from PLAZA for a given *M. truncatula* gene. Consequently, the number of CNS that had their closest genes evaluated varied from 2,600 to 3,805, depending on the method used to define the orthology (Table 2). The fraction of CNS which genes were considered orthologous ranged from 84.45% to 96.82% using Plaza’s Best-Hits-and-Inparalogs (BHI) family or Orthologous gene family methods, respectively (Table 2).

**Table 2.** Evaluation of orthology of the closest gene of each CNS specific to the NFC in *M. truncatula* and *G. max*, according to the orthology methods available on the PLAZA dicot v4.0 database.

| Method (Plaza DB) | Orthologous? |  |  |  |
| --- | --- | --- | --- | --- |
|  | No | Yes | Total | % |
| Tree-based ortholog | 293 | 2307 | 2600 | 88.74 |
| Orthologous gene family | 121 | 3,684 | 3,805 | 96.82 |
| Anchor point | 260 | 3158 | 3,418 | 92.39 |
| Best-Hits-and-Inparalogs (BHI) family | 590 | 3203 | 3,793 | 84.45 |

Aiming to characterize the largest number of CNS possible but also to focus on those most probable to have a biological effect, we opted to use the more inclusive “Orthologous gene family” method to filter the CNS-gene pairings. We ultimately identified 3,684 CNS mapped to orthologous genes for subsequent investigation (Supplemental File S2).

### CNS specific to the NFC clade

Considering the hypothesis that a predisposition or gain of the nodulation trait arose from a single event at the base of the NFC, we excluded all CNS detected in the NFC that were also observed within or overlapped in one or more nucleotides with a conserved region in the outgroup (593 CNS). Removal of these sequences resulted in 3,091 (83.90%) remaining for further analysis. The genomic coordinates of these CNS are available in the Supplemental File S3.

### Chromatin accessibility of CNS correlates with gene expression during nodule development

Chromatin accessibility of regulatory regions often contributes to modulate the expression of nearby genes. We previously generated global transcriptome and genome-wide chromatin accessibility data for *M. truncatula* (genotype Jemalong A17) roots, 0 min, 15 min, 30 min, 1 h, 2 h, 4 h, 8 h, and 24 h after treatment with rhizobium lipo-chitooligosaccharides (LCOs) (Knaack et al., 2021). Based on the ATAC-seq data, we observed that most of the remaining CNS (3,901 CNS, after filtering regions conserved in the outgroup) were in regions that display variation in chromatin accessibility following the LCO treatment (Supplemental Fig S3).

The detection of variation in the chromatin accessibility of CNS in response to the LCO treatment suggests a possible role of these regions in the transcriptional control of *Medicago* genes necessary for symbiotic signaling, rhizobial colonization, and nodule development. We therefore sought to identify CNS that could be important in these biological processes. The RNA-seq data collected from root samples was used to quantify transcription levels of the closest gene to each CNS classified as upstream, downstream, distal upstream, or distal downstream. In total, we tested 2,459 CNS and 1,155 genes for significant correlations between the chromatin accessibility profile of the respective CNS and a corresponding expression profile of a mapped gene, noting here that a single gene can be associated with multiple CNS. Based on Pearson’s correlation analysis, we detected 452 instances where the chromatin accessibility of a CNS is significatively correlated with the expression of the closest gene (p-value ≤ 0.05) responding to the LCO treatment. A total of 376 CNS correlated positively (ρ > 0.5; Figure 3) and 76 negatively (ρ < −0.5; Figure 4) with the expression profile of the closest gene. Considering genomic context, in 285 of the significantly correlated pairings the CNS are upstream of the gene (241 positively and 44 negatively correlated), 157 are downstream (125 positively and 32 negatively correlated) and 10 are distal upstream (all positively correlated). No significant correlation was observed for distal downstream CNS.

**Figure 3.**
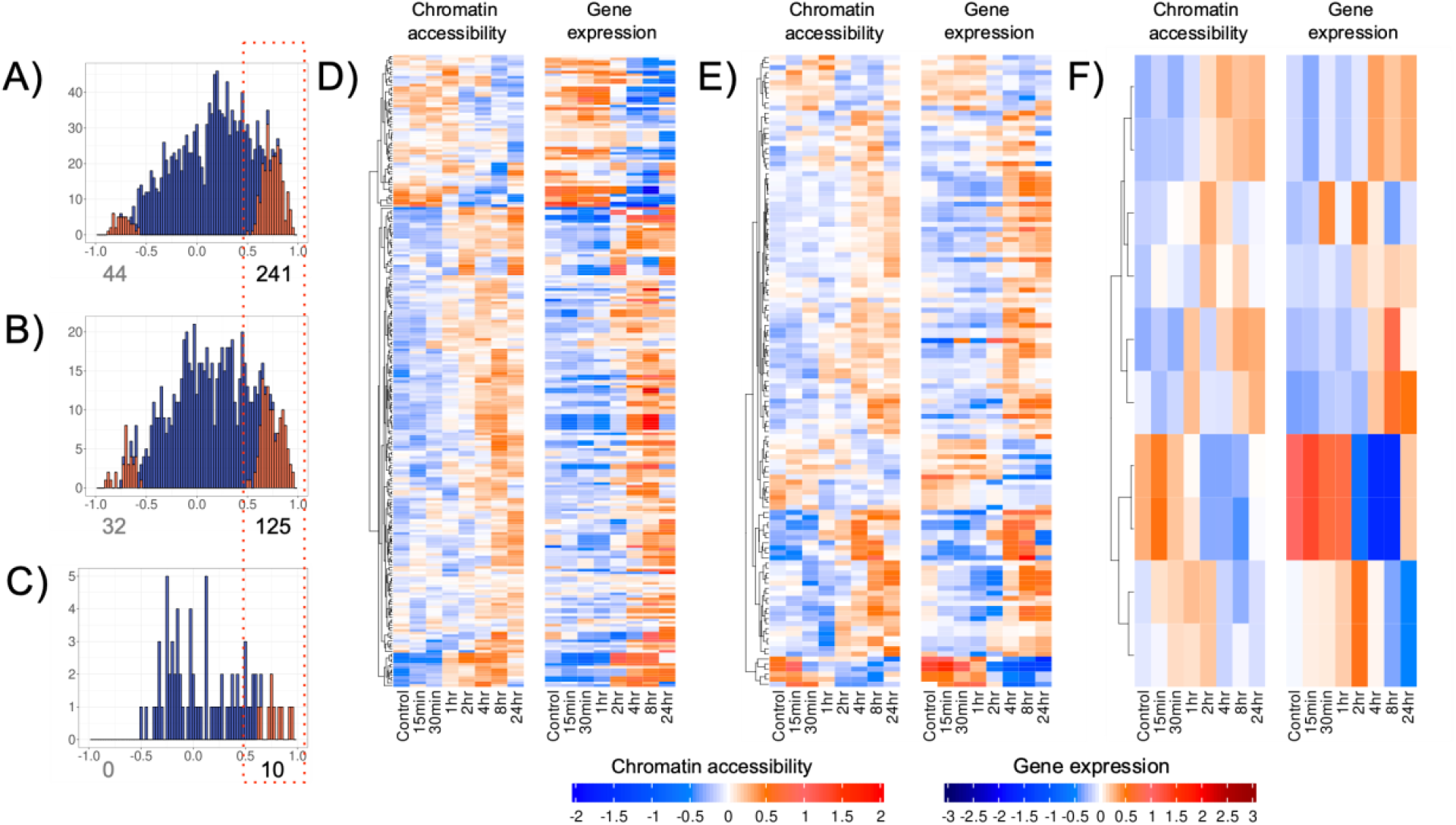
Chromatin accessibility of CNS and (mapped) gene expression profiles that are positively correlated. The profiles shown are based on CNS location relative to the closest mapped gene: A, D) CNS located upstream (0-2kb); B, E) downstream (0-1kb); and C, F distal upstream (2-10kb). The histograms (A, B and C) show the distribution of the Pearson correlation of CNS accessibility and gene expression profiles. The number of significant correlations (p-value < 0.05; highlighted in bold) is shown at the bottom of each histogram. For each pair of heatmaps (D, E and F) chromatin accessibility of the CNS (left), and the expression of the closest (mapped) gene (right) is shown. The rows of the heatmaps (both CNS accessibility and gene expression) were ordered by hierarchical clustering of the corresponding CNS accessibility profiles (note dendrograms, left of each heatmap). More than one CNS may be associated with a gene and the heatmaps consequently include repeated gene expression entries. No significant correlation was identified for the CNS classified as distal and downstream of the nearest gene.

**Figure 4.**
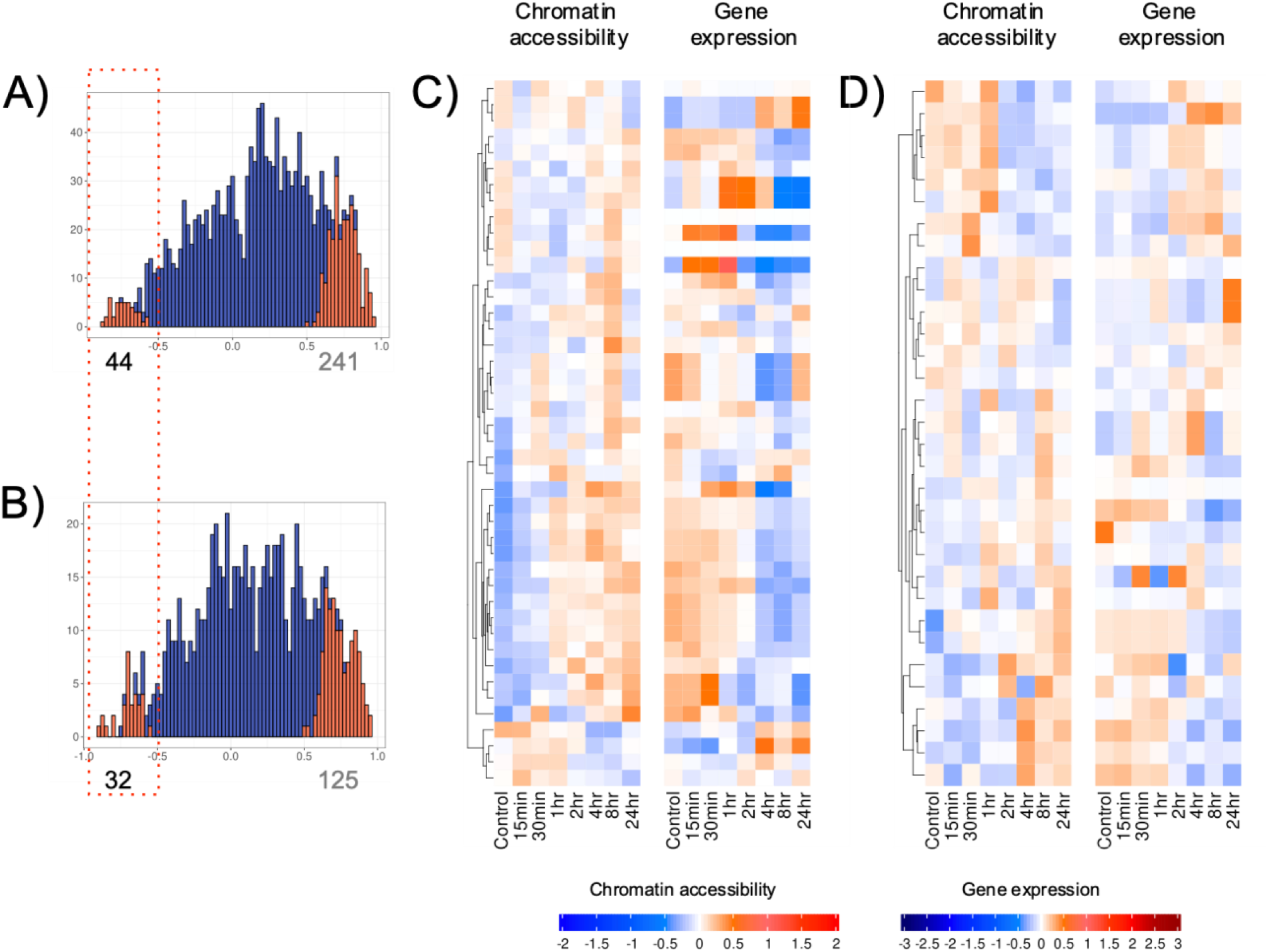
Chromatin accessibility and gene expression profiles of all CNS negatively correlated. The profiles are shown according to the CNS location from the closest gene, A, C) CNS located upstream (0-2kb); B, D) downstream (0-1kb). The histograms in A and B show the distribution of the correlation values. Significant correlations (p-value < 0.05, highlighted in bold) and the number of significant correlations is given beneath each histogram. For each pair of heatmaps (C and D), the profiles of the CNS chromatin accessibility (left) and expression of the closest gene are shown, where the rows of both are ordered by hierarchical clustering of the accessibility profiles (see dendrograms at left of heatmaps). Note that more than one CNS can be associated with a gene and consequently gene expression can have repeated entries. No significant correlation was identified for the CNS classified as distal (upstream or downstream).

### Conserved noncoding sequences associated with expression of nodulation genes

RNS is a complex developmental phenomenon in which a large number of genes are differentially regulated (Breakspear et al. 2014; Larrainzar et al. 2015; Jardinaud et al. 2016; Schiessl et al. 2019). These genes are collectively named as RNS genes and more than two hundred are known to date (Roy et al. 2020). From the 3,091 CNS specific to the NFC clade, we selected those whose closest gene is involved in RNS, and evaluated their chromatin accessibility and gene expression (Figure 5). A total of 38 CNS were located in proximity to 19 RNS genes. Ten CNS have a significant and positive correlation (p-value ≤ 0.05 and ρ > 0.5), and two have a significant and negative correlation (p-value ≤ 0.05 and ρ < −0.5).

**Figure 5.**
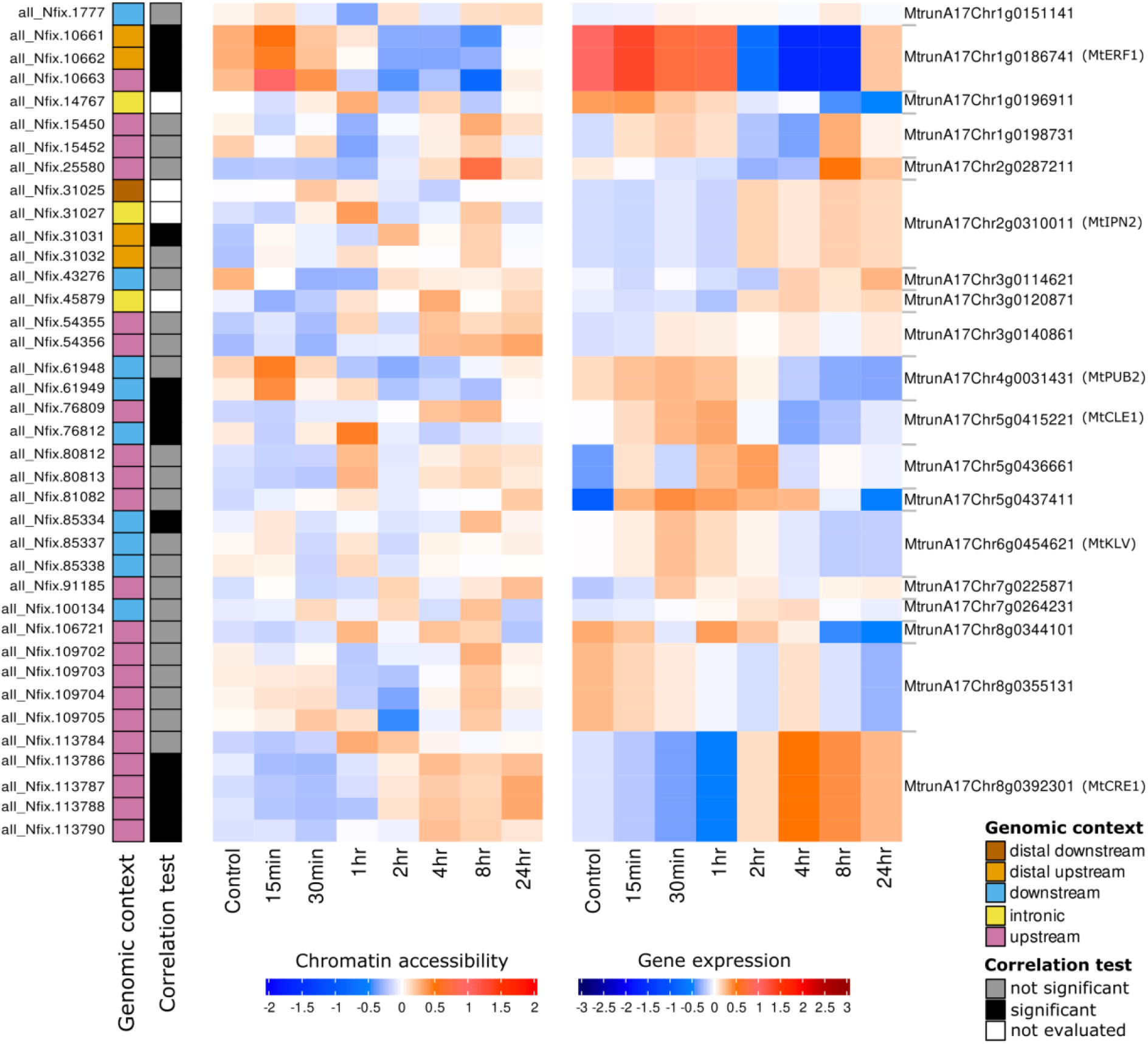
Chromatin accessibility profile of each CNS whose closest gene is an RNS gene (left) and their respective gene expression (right). The colors in the bar on the left indicate the genomic location of the CNS from the RNS gene (see legend). The second bar from the left indicates if the correlation is significant (black) or not significant (gray). For the intronic CNS regions, the correlation was not calculated (white). Common names are included for genes presenting significant correlation between their expression and the accessibility profile of at least one associated CNS. Only in the heatmaps of chromatin accessibility (left), the rows were hierarchically clustered. In the gene expression heatmaps, genes are organized in the same row as their associated CNS. Note that more than one CNS can be associated with a gene. Therefore, the heatmap representing gene expression has repeated entries.

The CNS significantly correlated with expression were associated with six RNS genes (Figure 5). These include three genes with well-established roles in nodulation (Figure 5): (1) *Cytokinin Response Element 1* (*MtCRE1*), encoding a histidine kinase cytokinin receptor is required for proper nodule organogenesis (Plet et al. 2011; Gonzalez-Rizzo et al. 2006); (2) *Interacting Protein of NSP2 (IPN2)*, encoding a member of MYB family transcription factor, activates the key nodulation gene *Nodule Inception (NIN)* in *Lotus japonicus* (Xiao et al. 2020); (3) *MtPUB2*, encoding an U-box (PUB)-type E3 ligase is involved in nodule homeostasis (Liu et al. 2018). We further observed association with three genes with potential roles in nodulation: (1) *MtCLE1* is a member of the Clavata3/ESR (CLE) gene family and is induced in nodules (Hastwell et al., 2017). Several CLE peptides are involved in the autoregulation of nodulation or regulation by nitrate or rhizobia (Nowak et al. 2019; Mens et al. 2021); (2) *MtERF1*, encoding an APETALA2/ethylene response factor (AP2/ERF) transcription factor (TF). *ERF Required for Nodulation 1* and *2* (*ERN1, ERN2*) play important roles during rhizobial infection (Cerri et al. 2012); (3) *MtKLV*, encoding a receptor-like kinase (*Klavier*). In *Lotus Japonicus*, *klavier* mediates the systemic negative regulation of nodulation and klavier mutants develop hypernodulation (Oka-Kira et al. 2005; Miyazawa et al. 2010). The two CNS with significant negative correlation were located upstream of the gene *MtCLE1* and downstream to the gene *MtKLV*, respectively.

Three of the 10 positively correlated CNS were located upstream (one) or distal upstream (two) the gene MtERF1 (ETHYLENE RESPONSE FACTOR 1; encodes an ethylene-responsive AP2 transcription factor). The CNS classified as upstream was located at the 5’ UTR while the two distal upstream CNS are located approximately five kilobases away from the gene. The two identified CNS might be part of one single conserved noncoding sequence. They are separated by only three nucleotides, which could be a consequence of misalignments or misclassification during the identification of conserved bases by PhastCons. Another distal-upstream CNS, located at approximately 8 kb of the gene MtIPN2, was identified (Figure 5). The remaining four significant and positively correlated CNS are located upstream of the gene *MtCRE1* (MtrunA17Chr8g0392301) (Figure 5).

Since CNS are known to harbor transcription factor binding motifs (TFBMs), we investigated if the twelve significantly correlated CNS contain TFBMs that could be implicated in the regulation of their corresponding RNS gene. PlantRegMap (Jin et al. 2017; Tian et al. 2020) was used to search for the presence of TFBMs, and 41 TFBMs were identified in six of the CNS. In some cases, all TFBMs identified in a CNS belong to the same family of TFs. In contrast, in other CNS, TFBMs of multiple families of TFs were identified (Supplemental File S4).

### Validation of a CNS associated with the function of *MtCRE1* in nodulation

Due to its role as a central regulator of nodule organogenesis and the availability of *Mtcre1-1* mutants in the genotype background (Jemalong A17) used in this study (Plet et al. 2011), it represents a compelling candidate for the experimental investigation of CNS’ role in the regulation of genes related to nitrogen fixation. To test this hypothesis, we evaluated if the deletion of the four CNS associated with *MtCRE1* would hinder the occurrence of RNS. These CNS were in the 5’ UTR region, where one was in the 5’ UTR and the other three were in an intron that divides the UTR into two segments. We notice that an additional CNS located in the 5’ UTR was removed during the filtering steps because it was not possible to recover its corresponding coordinates in the *G. max* genome. Considering that all other CNS located in *MtCRE1* aligned to an orthologous gene in *G. max* (LOC100789894; also known as Glyma.05G241600), we considered the absence of alignment to this region of the *G. max* genome a possible error and included it in the experimental validation (Supplemental Fig S4). To investigate if the five CNS identified in *MtCRE1* are required for nodulation, three versions of *MtCRE1*’s promoter containing deletions of the distal two CNS (Δ2CNS), or proximal three CNS (Δ3CNS), or all CNS (Δ5CNS) were engineered.

Composite *Medicago truncatula* plants in the *Mtcre1-1* background were generated by the transformation of roots with a construct containing *MtCRE1* either under the wild-type promoter (positive control) or one of the three engineered promoters. In addition, a construct lacking the *MtCRE1* cassette was used as an empty vector (negative control). The transformed roots were inoculated with *S. meliloti* constitutively expressing *lacZ*, which allows the identification of successful infection. Two weeks post inoculation, a gradual decrease in the number of nodules in the plants containing the CNS deletions was observed (Figure 6). Statistical analysis shows a significant reduction in the number of nodules in the comparison between the wild-type and Δ5CNS roots, providing evidence that the CNS are required for nodule organogenesis. Moreover, while the differences were not statistically significant, Δ2CNS and Δ3CNS showed a reduced number of nodules, suggesting that deletions of these CNS may partially impair the capacity of the plants to engage in RNS.

**Figure 6.**
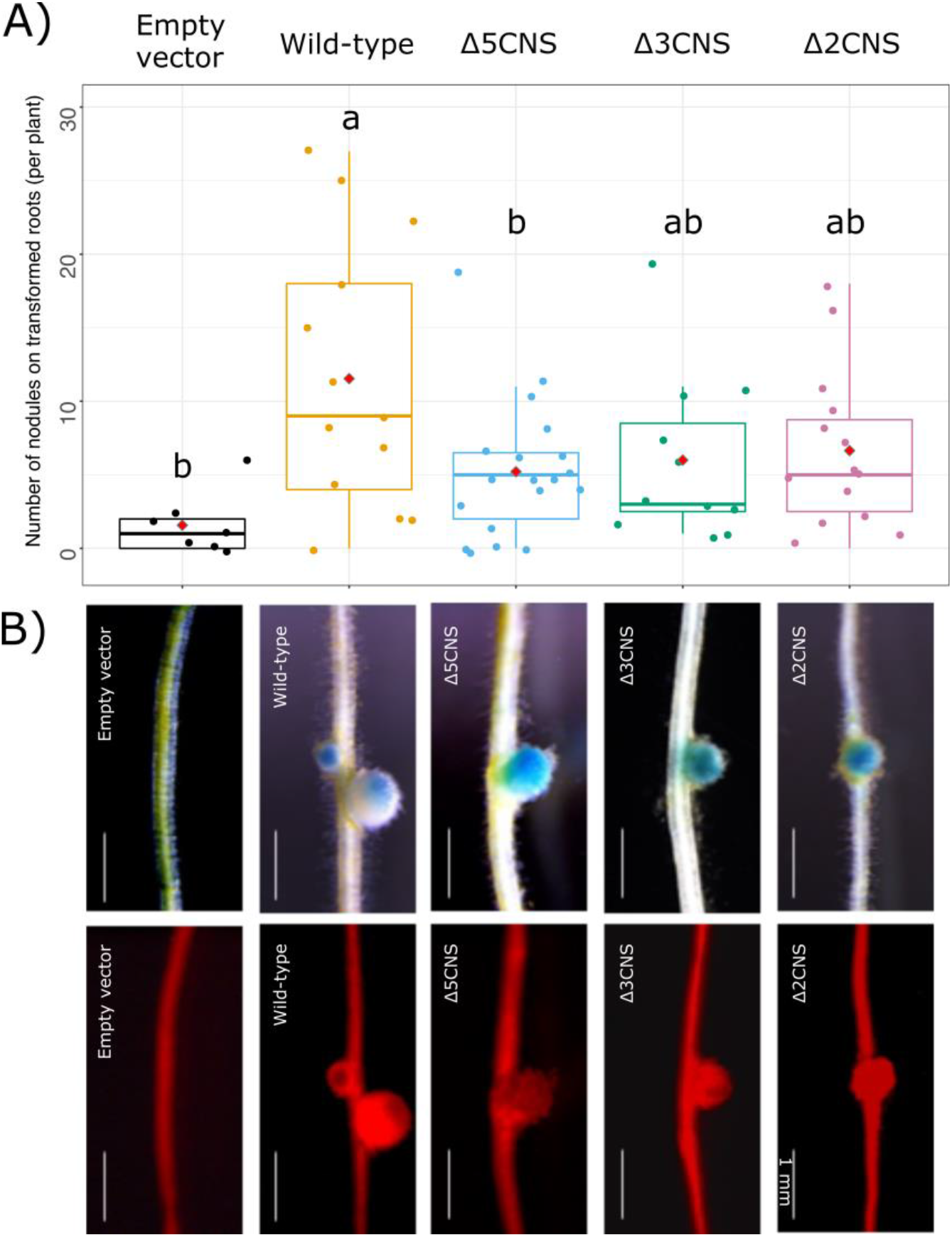
Role of the *CRE1* CNS in nodulation. Radicles of *M. truncatula* Jemalong A17 were inoculated with *Agrobacterium rhizogenes* MSU440, expressing *MtCRE1* either under the native (WT) promoter or one of the constructions containing two, three or all five CNS related to this gene deleted (Δ2CNS, Δ3CNS, and Δ5CNS, respectively). In addition, an empty vector was also used as a control. Three weeks after transformation, the roots of the composite plants were screened for red fluorescence of tdTomato on the binary vector. Plants with transformed roots were transferred to growth pouches and acclimated for a week, and then inoculated with *S. meliloti 1021* harboring pXLGD4 constitutively expressing *lacZ.* Two weeks after inoculation, live seedlings were stained with X-gal, the nodules scored and imaged. In A), it is shown the distribution of the total number of nodules on transformed roots per plant. Columns not connected by the same letter(s) are significantly different at *α* = 0.05, ANOVA followed by Tukey’s post-hoc test for multiple comparisons. Data from three independent rounds of transformation with *n* = 7-19 composite plant per genotype. Red diamantes show the average. In B), are shown representative X-gal-stained nodules of the transformed roots visualized in the brightfield (upper panel) and tdTomato fluorescence (lower panel). Scale bar = 1 mm.

## DISCUSSION

Significant progress has been achieved in uncovering the genomic elements underpinning the capacity of plants to engage in RNS, with more than two hundred essential genes identified to date (Roy et al. 2020). Nonetheless, the current knowledge has not enabled the long-standing goal of transferring the genetic toolkit required for RNS to plant species lacking this capability. Moreover, comparative genomics indicates that species lacking RNS (including those outside the NFC) contain the majority or all of the required genes (Griesmann et al. 2018). However, the gain of a trait can also be driven by the differential regulation of a similar gene ensemble. Thus, the evolution of regulatory elements may have been critical for the appearance and maintenance of RNS in the NFC.

With few exceptions (Liu et al. 2019), the potential role of regulatory elements in RNS has remained largely unexplored. Here, we applied comparative genomics to identify thousands of CNS in nodulating species of the NFC clade. Moreover, using chromatin accessibility and RNA sequencing data, we defined the CNS most prone to affect the function of genes essential to RNS. We select 34 genomes that were used for multiple whole-genome alignments (WGA), generating two groups: NFC, containing 25 species, and the outgroup, containing nine species. Some issues can emerge from the usage of whole-genome alignments. Especially in plant species, where genome duplications are common, and a large portion of the genome corresponds to repetitive elements, identifying the correct alignments can be challenging. We deployed a workflow previously used to detect CNS in grasses (CNSpipeline; Liang *et al*., 2018). As part of this workflow, split-last (Frith and Kawaguchi 2015) was applied to identify the correct alignment for each region. In our data, the genome comparison with *G. max* showed that, for most CNS, the alignment was established with the right orthologous region. Most CNS were classified in the same genome context in both *M. truncatula* and *G. max* (Figure 2).

Chromatin accessibility in noncoding regions of the genome is often used as a proxy to identify *cis*-regulatory elements (de Velde et al. 2014; Lu et al. 2019; Song et al. 2021; Marand et al. 2021) since these regions tend to be enriched in TFBMs. We demonstrated that the chromatin accessibility profile (measured by ATAC-seq) of hundreds of CNS strongly correlates with the gene expression of their closest gene during the first 24 hours of Medicago’s response to LCO stimulus. While the CNS with significant correlation represents only a fraction of the set of NFC-specific CNS, the remaining CNS cannot be disqualified as regulatory elements involved in the RNS. Those regions could be related to the regulation of genes in which the response is triggered at different stages of the response to the stimulus. Alternatively, they may regulate genes required for the rhizobium infection that are not responsive to LCO treatment. Therefore, studies including more time-points and following the plant response to the rhizobium infection are still necessary to thoroughly evaluate these CNS.

It is worth mentioning that, by considering only the closest gene of each CNS, we may have failed to detect significant associations between the chromatin accessibility of a CNS and its target gene because a CNS could act on the expression of a distal gene or multiple nearby genes. Nonetheless, we identified several significant correlations between CNS’s chromatin accessibility and the expression of genes essential for the RNS, pointing to functional roles for these elements (Figure 5). We validated the function of five CNS located upstream of the gene *MtCRE1*, four of which showed a significant correlation between their chromatin accessibility and the gene expression. The deletion of these regions produced fewer nodules on *M. truncatula* after the infection by *S. meliloti* (Figure 6) demonstrating that these CNS are required for the correct functioning of *MtCRE1* during RNS. The observed effect may be associated in part with the disruption of the 5’ UTR region of *MtCRE1*.

A complete understanding of the genomic elements required for a plant to engage in RNS will enable engineering this phenotype in plants outside the NFC. Achieving this goal will require the evaluation of genomes beyond the coding regions, to capture the elements involved in regulating essential genes and that are in the noncoding segments of the genome. In this work, we compared the genome of species in the NFC and identified hundreds of CNS in *M. truncatula* that potentially affect RNS. Moreover, we show experimental evidence that CNS are required for the correct functioning of *MtCRE1* and for the establishment of RNS. To the best of our knowledge, this is the first genome-wide study attempting to connect conserved noncoding sequences to RNS, providing a foundation for uncovering essential CNS sites in this process.

## METHODS

### Species selection and genome sequence quality control

We searched the National Center for Biotechnology Information (NCBI) RefSeq and GenBank databases for all publicly available genomes belonging to orders containing species capable of nitrogen fixation (Fabales, Fagales, Cucurbitales, and Rosales), as of October 2018. For some species, an improved version of the genome assembly available in Phytozome (Goodstein et al. 2012) was used instead. Because we targeted the identification of conserved regions among plant species capable of engaging in RNS, we discarded those unable to associate with nitrogen-fixing bacteria based on existing information in the literature. Moreover, we selected a group of species not capable of engaging in RNS, belonging to orders outside the NFC, but phylogenetically near this clade (outgroup). While these orders are outside of the NFC, their phylogenetic proximity to the clade implies lower overall sequence divergence, simplifying the detection of functionally relevant differences.

To verify the quality of these genomes, we applied two analyses. The assembly-stats v.1.0.1 algorithm (https://github.com/sanger-pathogens/assembly-stats) was implemented to investigate the contiguity of the assemblies, and BUSCO v.3.02 (Seppey et al. 2019) was used to evaluate the completeness of the corresponding gene annotations based on the embryophyta_odb10 dataset (https://busco-archive.ezlab.org/v3/frame_wget.html). We constrained all analyses to species for which the assembled genome contained at least 80% of the BUSCO genes in this dataset.

### Whole-genome alignments

Multiple sequence alignments of whole genomes were generated for each of the two sets of selected genomes (NFC and outgroup) separately. A previously developed workflow was implemented (Liang et al. 2018) to build these multiple alignments. First, each genome was submitted to a pairwise whole-genome alignment to the genome of a reference species using LAST v916 (Kiełbasa et al. 2011). The *M. truncatula* (v.5; Pecrix *et al*., 2018) genome was used as the reference. Before the alignment procedure, simple repetitive elements were masked from all genome sequences using the algorithm tantan (Frith 2011b, 2011a). The alignments were generated using the lastdb argument −uMAM8, and lastal arguments −p HOXD70 −e 4000 −C 2 −m 100. Next, the split-last algorithm (Frith and Kawaguchi 2015) was used to improve the selection of orthologous sequences. Finally, the pairwise whole-genome alignments were assembled using axtChain and chainNet (Kent et al. 2003).

All pairwise alignments were combined in a multiple-alignments file using ROAST (reference-dependent multiple alignment tool), which merges the alignments using a phylogenetic tree-guided approach (http://www.bx.psu.edu/~cathy/toast-roast.tmp/README.toast-roast.html, last access in Jun 2021). The same procedure was employed to generate the whole-genome alignment of the two groups described above (NFC and outgroup), including the use of *M. truncatula* as the reference. The phylogenetic tree used to guide ROAST is shown in Figure 1.

### Identification of conserved noncoding regions in species capable of engaging in RNS

PhastCons (Siepel et al. 2005) was used to estimate the conservation score of the *M. truncatula* genome regions contained within multiple alignments. Briefly, for each multiple genome alignment group (NFC and outgroup), we first applied phyloFit from the PHAST toolkit, version 1.5, to estimate a background model of sequence evolution. Next, we used the phastCons estimate-tree function to learn models of conserved and non-conserved sequence evolution, respectively. Finally, the phastCons most-conserved function was applied on each chromosome at a time to identify conserved and diverged sequences.

The regions identified were further filtered to exclude those: (1) smaller than five nucleotides, (2) overlapping (>=1 bp) transposable elements and noncoding RNAs, and (3) contained within the mitochondrial or chloroplast genomes. Because this analysis focused on identifying putative regulatory sequences associated with the evolution of RNS, we also excluded from further investigation the regions overlapping in one or more bases with regions annotated as coding sequences in the *M. truncatula* genome. The remaining conserved regions were compared with a subset of the non-redundant protein database (nr), containing all *viridiplantae* species, using Blastx v,2.10.1+. All regions with an e-value <= 0.01 were excluded. This step aims to remove coding regions that might exist but were not annotated in the *M. truncatula* genome version 5. The analysis workflow for the definition of the CNS regions is shown in Supplemental Fig S5.

### Genomic context of CNS and ortholog comparison with Glycine max

To perform filtering of the CNS based on synteny to other species, they were classified in six categories according to their distance to the closest gene in the *M. truncatula* genome. The categories were (1) intronic (locate inside an intron of a gene), (2) downstream (up to 2 kb downstream of the translation stop site), (3) upstream (up to 2 kb upstream of the translation start site), (4) distal upstream (2-10 kb upstream of the translation start site), (5) distal downstream (2-10 kb downstream of the translation stop site), and (6) intergenic (all CNS not classified in the previous five categories). We considered the translation start site and translation stop site as the reference coordinates to permit the comparison with the soybean (*G. max*) genome, for which UTR coordinates are not described in the NCBI’s annotation. The CNS located in the *M. truncatula* UTRs’ were classified as downstream (3’ UTR) or upstream (5’ UTR). Using the coordinates of the genome alignment, we located the CNS in the *G. max* genome and classified them following the same approach.

Next, we examined if the CNS identified in the *M. truncatula* genome were in orthologous regions of *G. max*; v.2.1 (Schmutz et al. 2010). The soybean genome was selected for this analysis because gene orthology to *M. truncatula* is well-established and readily available in the PLAZA database (Van Bel et al. 2018). Moreover, for the *G. max* genome, it was possible to recover the relationship between the NCBI’s (Entrez) locus tags and the identification adopted in the PLAZA database, allowing the comparison. Because PLAZA uses the gene identification of version 4 of the *M. truncatula* genome assembly and annotation, we used the information from the genome portal of *M. truncatula* assembly version 5 (https://medicago.toulouse.inra.fr/MtrunA17r5.0-ANR/, last accessed in Jun 2021) to recover the gene identification equivalent in the newer version (v5). In some cases, two or more genes identified in v4 corresponded to a single gene in v5. We considered a gene an orthologous if at least one of the v4 gene identifications was deemed orthologous of the corresponding gene in *G. max*. Only CNS where the closest gene was considered an ortholog were kept for further evaluation.

### Chromatin accessibility of CNS and gene expression of associated genes

To identify CNS that are associated with regulatory changes triggered by rhizobial infection and nodule formation, we examined chromatin accessibility (ATAC-seq) and gene expression (RNA-seq) data captured after *M. truncatula* root treatment LCOs from *S. meliloti* 2011 (Knaack et al., 2021). Briefly, for generating this data, roots from wild-type (WT) seedling (reference accession Jemalong A17) were immersed in a solution of purified LCOs derived from *S. meliloti* or 0.005% ethanol solution (control) for 1 h and collected posteriorly at specific intervals (0 h, 15 min, 30 min, 1 h, 2 h, 4 h, 8 h, and 24 h). We obtained the quantile-normalized and log-transformed TPM (Transcripts Per Kilobase Million) expression values for all *M. truncatula* genes (Knaack et al., 2021). Additionally, ATAC-seq read counts were aggregated and normalized for each CNS. Specifically, for each CNS the mean per-bp coverage was obtained and log-ratio-transformed relative to genome-wide mean coverage for each time-point, producing a normalized accessibility profile across time. Quantile normalization was subsequently applied across the time course to the log-ratio-normalized values.

To evaluate the relationship between gene expression and CNS accessibility, we carried out a correlation analysis, as described previously (Knaack et al., 2021). Briefly, we first performed a zero-mean transformation of each gene’s expression profile and CNS’s accessibility profiles. Pearson’s correlation (ρ) between the accessibility profiles of each site and the expression profile of the closest gene across all eight time-points was calculated. In this step, only the CNS classified as downstream, upstream, distal upstream, or distal downstream were included. To assess the significance of the relationship between gene expression and chromatin accessibility of the CNS, a null distribution of correlations from 1,000 random permutations of the chromatin accessibility score and the gene expression in the eight time points was generated. We computed a p-value that estimates the probability of observing a correlation in the permuted data higher than the observed correlation, treating positive and negative correlation separately. In practice a p-value threshold of ≤ 0.05 for significant correlation was used. Also, we focus on the CNS which chromatin accessibility and gene expression produced strong correlations (ρ > 0.50 or < −0.50).

### Selection of CNS potentially involved in nitrogen fixation

To exclude CNS present in species outside the NFC, we removed those that overlap (>=1 bp) with CNS regions called for the outgroup species. The remaining filtered CNS consists of those with a potential role in RNS because of their conservation within this clade.

A large number of genes have been previously implicated in the RNS, collectively known as RNS genes. These genes were identified based on the current literature in *M. truncatula* symbiosis and orthology with other species within the NFC. To determine CNS that may contribute to the regulation of these genes, we initially identified those conserved regions classified as upstream (0-2kb), downstream (0-2kb), distal upstream (2-10kb), or distal downstream (2-10kb), of the RNS genes. We further selected those regions where a significant correlation between chromatin accessibility and gene expression was detected. For the selected CNS, a search for transcription factor binding motifs (TFBMs) was conducted using the PlantRegMap database (Tian et al. 2020; Jin et al. 2017).

### Experimental validation of CNS associated with the R gene *MtCRE1*

To evaluate the functional role of CNS identified upstream of the *MtCRE1* gene, we used the Golden Gate modular cloning system to transform *Mtcre1-1* mutant (Plet et al., 2010) and insert a functional copy of *MtCRE1* fused with three different versions of the 2.5 kb region upstream the translation starting point, containing the promoter region and the 5’ UTR, synthesized by Synbio Technologies (NJ, USA). In each version, the sequences corresponding to two or more of the five CNS identified in the 5’ UTR of *MtCRE1* were deleted, as following. In the first version, CAGATCAAGA (−807 to −797; CNS = all_Nfix.113790) and GTGTTCATTGTA (−767 to −765; CNS = all_Nfix.113790) were deleted (Δ2CNS), in the second version GGTCAATGTTTGGTTTTTTTGATTAAAC (−628 to −600; CNS = all_Nfix.113788), TGAACGTGCTTTTGTTTGTCCT (−589 to −567; CNS = all_Nfix.113787), and TTCACGCCTAGAGAGCATGATTAGA (−552 to −526; CNS = all_Nfix.113786) were deleted (Δ3CNS), and in the third version all five regions were deleted (Δ5CNS), see Supplemental FigS4. The two CNS deleted in Δ2CNS are in the 5’ UTR, while the three CNS deleted in Δ3CNS are in an intron that splits the 5’ UTR (based on Jemalong A17 genome v.5.0; Supplemental Fig S4.). Golden Gate cloning was utilized as previously described (Engler et al. 2014).

To assemble the transcriptional units, level 0 parts of each version of the promoter were created with *MtCRE1* coding region and 35S terminator (MoClo Plant Parts Kit-pICH41414) into the MoClo toolkit level 1 acceptor-position2-reverse (pICH47811) vector using the Golden Gate BsaI/T4 restriction ligation reaction (Engler et al. 2014; Weber et al. 2011). Finally, each Level 1 transcriptional unit containing a different version of the promoter were assembled into the MoClo Level 2 acceptor (pAGM4673) together with the Level 1 transcriptional units containing the fluorescent marker of transformation, p35S::tdTOMATO-ER::tNOS, using the Level 2 BpI/T4 restriction ligation reaction (Weber et al. 2011, Engler et al. 2014).

### Generation of composite plants and nodulation assay

*Mtcre1* mutant or their wild-type sibling radicles were inoculated with *Agrobacterium rhizogenes* MSU440 as previously described (Boisson-Dernier et al. 2001). Three weeks after transformation with MSU440, the roots were screened for red fluorescence of *tdTomato* on the binary vector. The composite plants with red fluorescent roots were transferred to growth pouches (https://mega-international.com/tech-info/) containing Modified Nodulation Medium (Chakraborty et al. 2021). The plants were acclimated for a week and then inoculated with *S. meliloti* 1021, harboring pXLGD4 and expressing *lacZ* under the *hemA* promoter (Leong et al. 1985). The nutrient medium was replenished as required. Two weeks after inoculation, live seedlings were stained for *lacZ* (5 mM potassium ferrocyanide, 5 mM potassium ferricyanide, and 0.08% X-gal in 0.1 M PIPES, pH 7) overnight at 37°C. Roots were rinsed in distilled water and observed under Leica fluorescence stereomicroscope. Nodule presence was scored at 14 days post-inoculation.

## DATA ACCESS

All genomes utilized in this study are available via NCBI and or Phytozome (see Supplemental File 1 for instructions of how to download them). The code to reproduce the analysis is also freely available and can be found on GitHub (https://github.com/KirstLab/CNS_Nitrogen_Fixing_Clade).

Both RNA-seq and ATAC-seq data used in this publication were generated and published by Knaack et al., 2021. The data have also been deposited in NCBI’s Gene Expression Omnibus (https://www.ncbi.nlm.nih.gov/geo/) and are accessible through accession number GSE154845 (https://www.ncbi.nlm.nih.gov/geo/query/acc.cgi?acc=GSE154845).

## ACKNOWLEDGMENTS

We thank the US Department of Energy, Office of Science Biological and Environmental Research for funding this study (DE-SC0018247 to M.K., S.R. and J.M.A).

