## Supplemental Fig S1 for "Functional and comparative genomics reveals conserved noncoding sequences in the nitrogen-fixing clade"

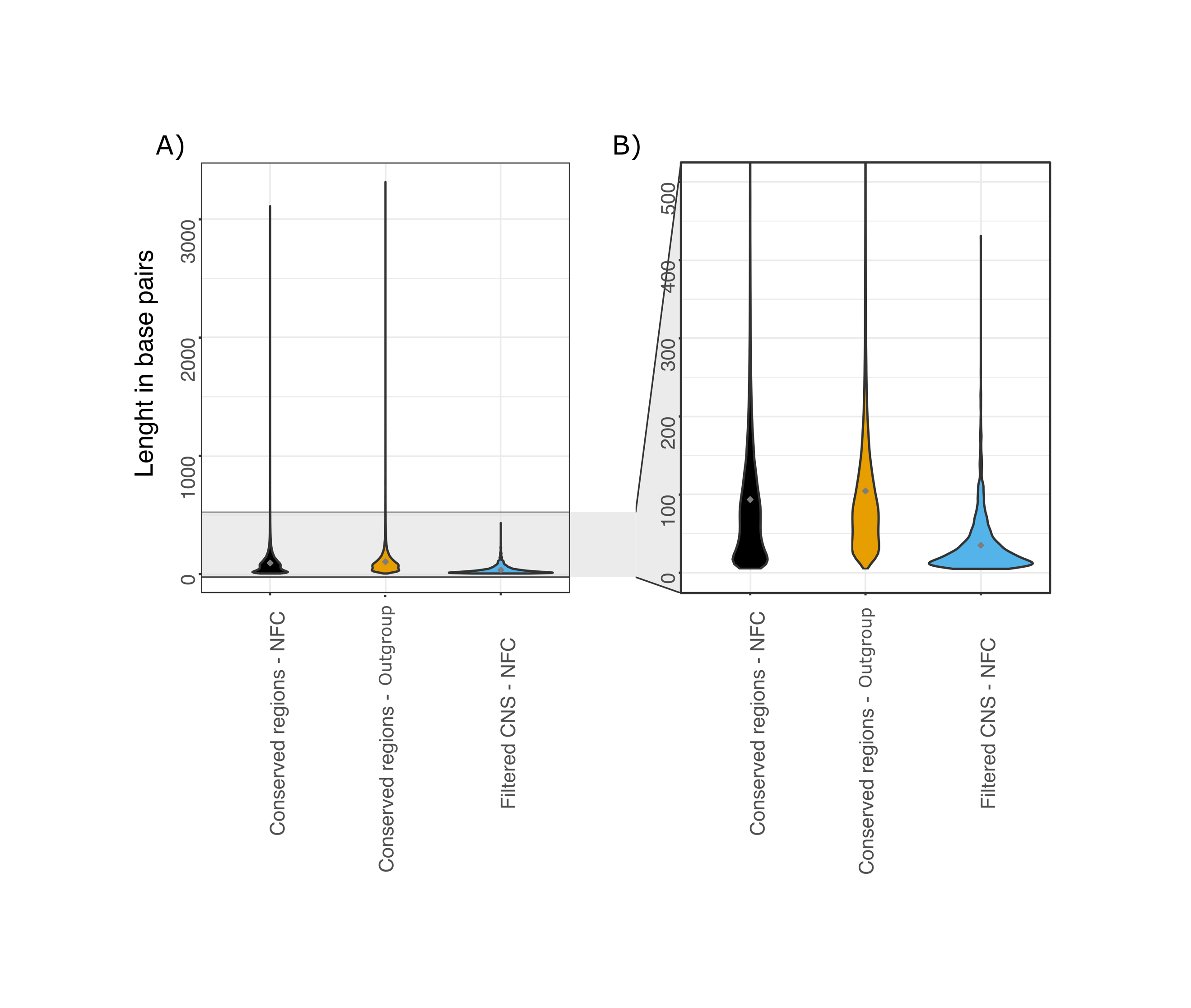

**Supplemental Figure S1. CNS specific of the nitrogen-fixing clade are in general smaller than the conserved regions of the genome.** A) Violin plot showing the distribution of the regions’ length, including the outliers. B) Violin plot showing the distribution of the regions’ length with emphasis in the length range that contains most of the regions, therefore, excluding the outliers. In black, are shown the conserved regions identified in the nitrogen-fixing clade and, in yellow, the conserved regions of the outgroup. In blue, is show the set of conserved noncoding sequences specifically detected in the NFC (after the exclusion of CNS that overlap with coding sequences or other regions that are conserved in the outgroup).
