## Supplemental Fig S2 for "Functional and comparative genomics reveals conserved noncoding sequences in the nitrogen-fixing clade"

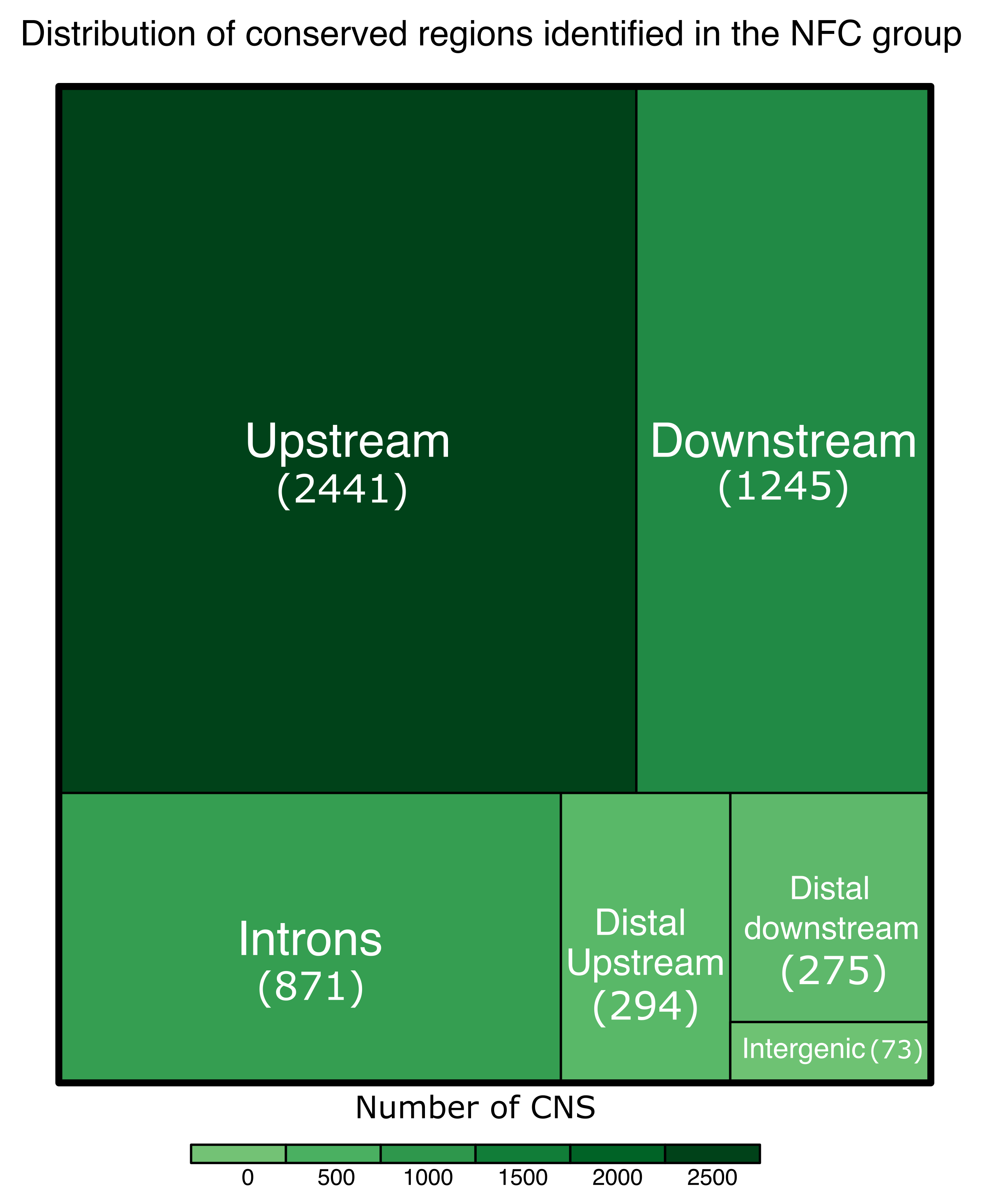

**Supplemental Figure S2. Distribution of the CNS identified on the nitrogen-fixing clade according to their genome context.** The size of the rectangle, and its color shade, reflect the proportion of CNS in each category.
