## Supplemental Fig S3 for "Functional and comparative genomics reveals conserved noncoding sequences in the nitrogen-fixing clade"

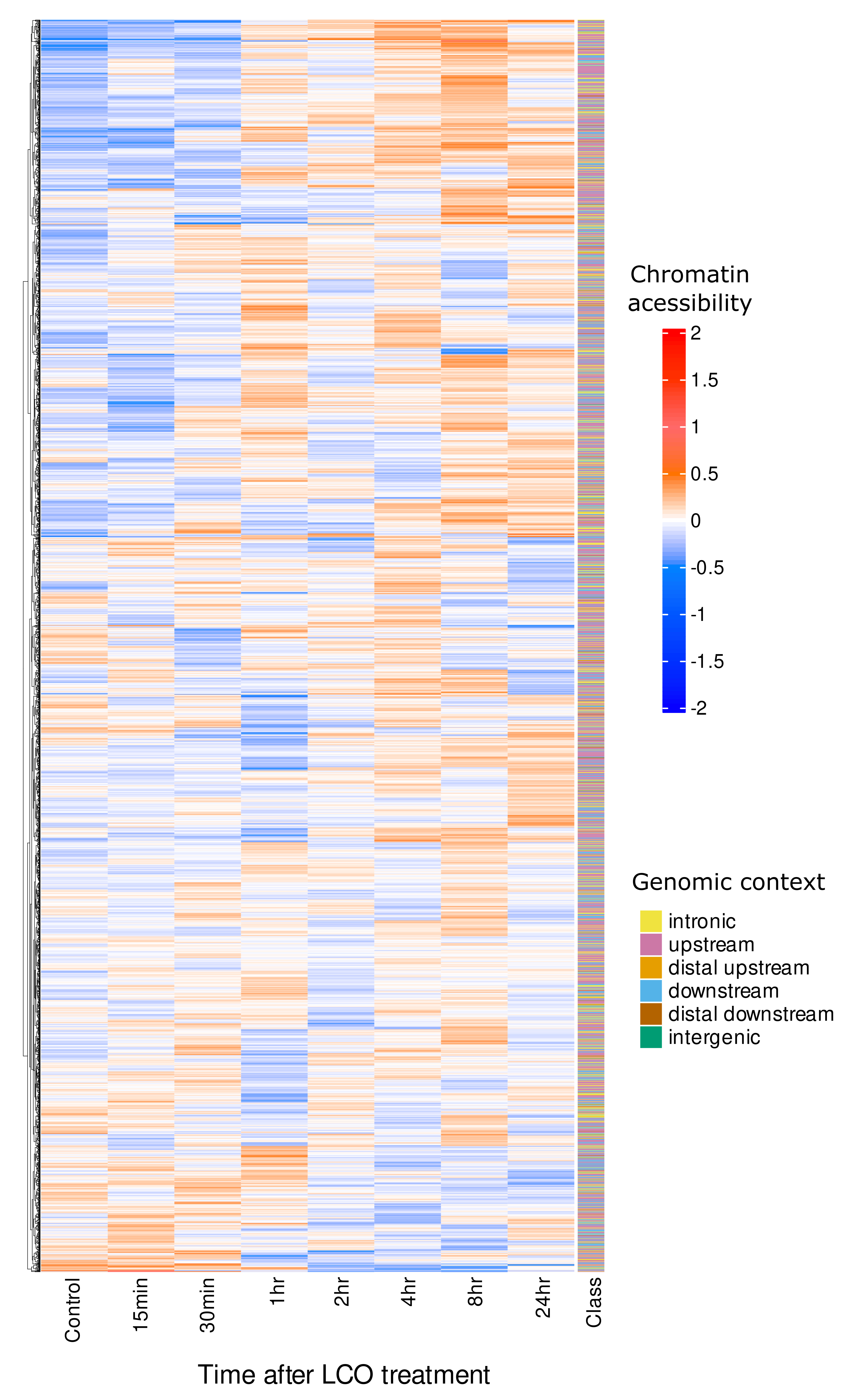

**Supplemental Figure S3.** Chromatin accessibility profile of the CNS in response to LCO treatment as measured by ATAC-seq in a time-series experiment. All 5,199 CNS identified for the NFN clade are represented. The x-axis represents the time after the exposure to the treatment in chronological order. In the y axis, the CNS are ordered according to their accessibility profile by a hierarchical clustering algorithm. In the column to the right, the classification of the CNS is shown according to its distance to the closest gene.
