## Supplemental Fig S4 for "Functional and comparative genomics reveals conserved noncoding sequences in the nitrogen-fixing clade"

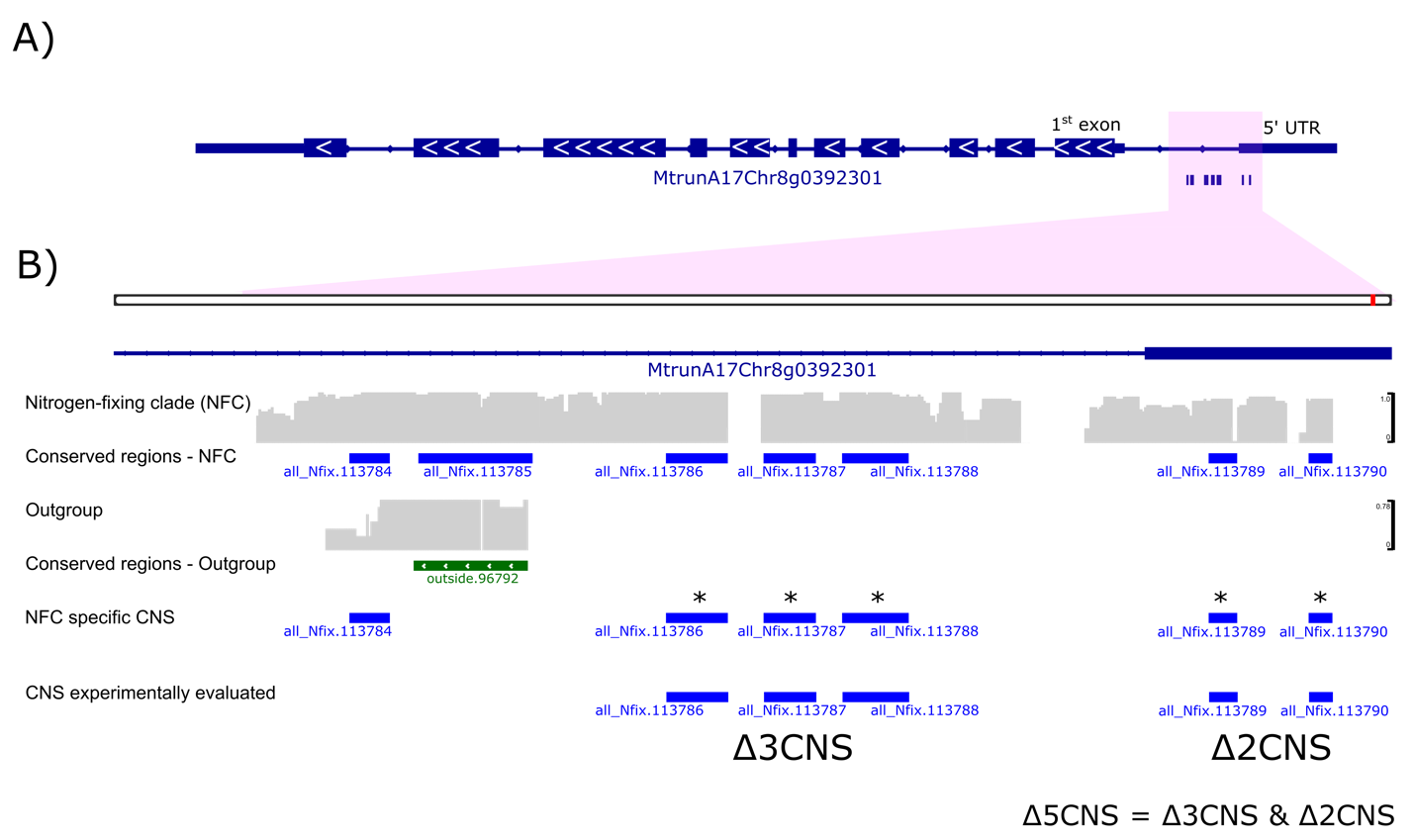

**Supplemental Fig S4.** Conserved noncoding regions in the gene *MtCRE1*. A) Gene model of *MtCRE1.* In blue, seven CNS detected upstream of the translation starting site are shown. B) Expanded representation of the region containing the detected CNS. Five of the seven CNS have a significant correlation between chromatin accessibility and gene expression of *MtCRE1* (Figure 5) and are marked with an asterisk. The remaining two CNS, all_Nfix.113784 and all_Nfix.113785, were excluded from further analysis because they were detected in the outgroup. In gray, the coverage tracks of the whole-genome alignment of species in the NFC and outgroup are shown. The region that is conserved in the NFC and the outgroup is shown in green.
