## Supplemental Fig S5 for "Functional and comparative genomics reveals conserved noncoding sequences in the nitrogen-fixing clade"

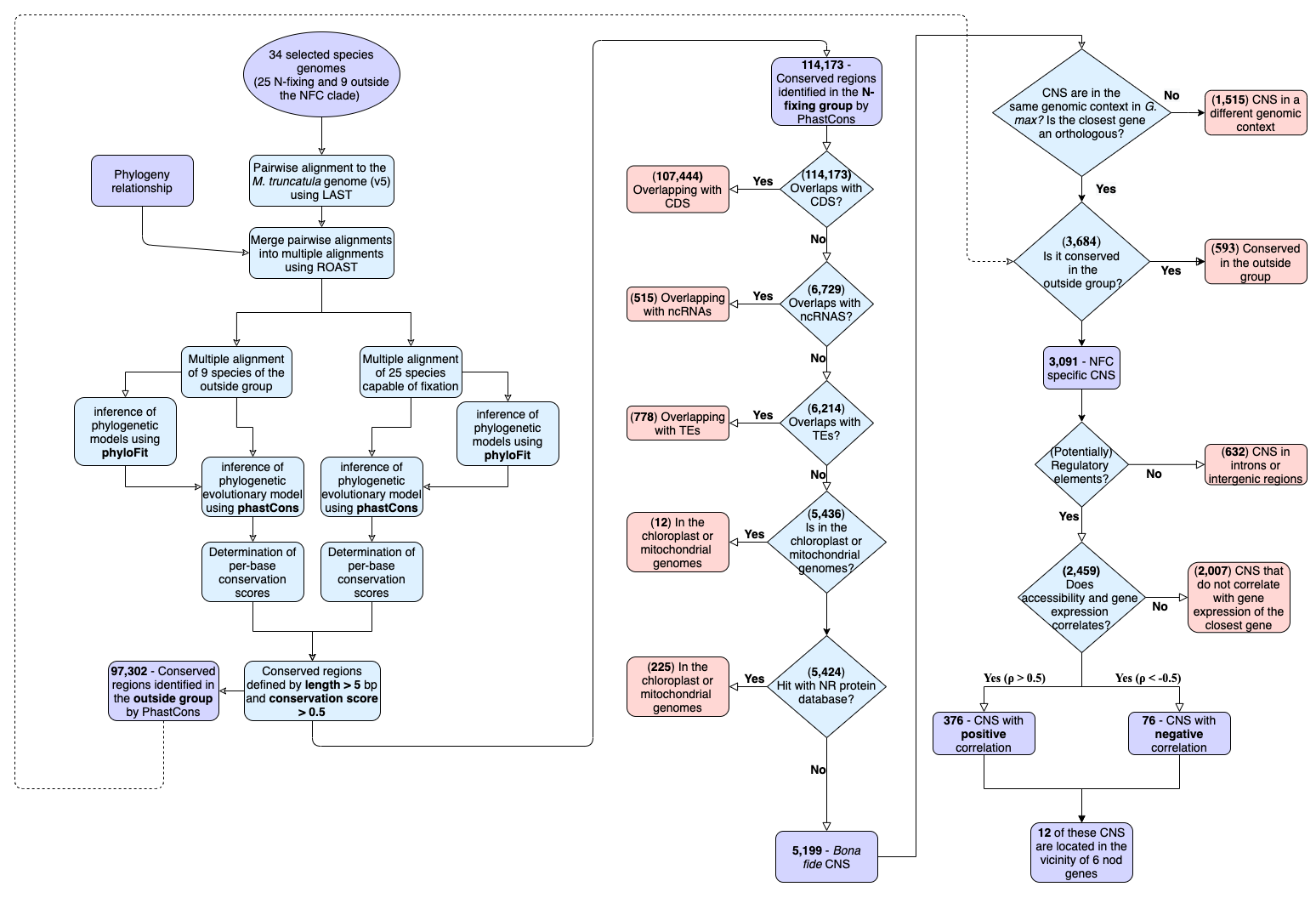

**Supplemental Fig S5.** Workflow representing the main analytical steps to generate the alignments, define the conserved noncoding sequences (CNS), and select the CNS that are potentially related to root nodule symbiosis.
